# Development of a yeast whole-cell biocatalyst for MHET conversion into terephthalic acid and ethylene glycol

**DOI:** 10.1101/2022.10.30.514423

**Authors:** Raphael Loll-Krippleber, Victoria Sajtovich, Michael W. Ferguson, Brandon Ho, Brandon J. Payliss, Joseph Bellissimo, Sydney Peters, Haley D. M. Wyatt, Grant W. Brown

## Abstract

**Background:** Over the 70 years since the introduction of plastic into everyday items, plastic waste has become an increasing problem. With over 360 million tonnes of plastics produced every year, solutions for plastic recycling and plastic waste reduction are sorely needed. Recently, multiple enzymes capable of degrading PET (polyethylene teraphthalate) plastic have been identified and engineered. In particular, the enzymes PETase and MHETase from *Ideonella sakaiensis* depolymerize PET into the two building blocks used for its synthesis, ethylene glycol (EG) and terephthalic acid (TPA). Importantly, EG and TPA can be re-used for PET synthesis allowing complete and sustainable PET recycling.

**Results:** In this study, we used *Saccharomyces cerevisiae* as a platform to develop a whole-cell catalyst expressing the MHETase enzyme, which converts MHET (monohydroxyethyl terephthalate) into TPA and EG. We assessed six expression architectures and identified those resulting in efficient MHETase expression on the yeast cell surface. We show that the MHETase whole-cell catalyst has activity comparable to recombinant MHETase purified from *Escherichia coli*. Finally, we demonstrate that surface displayed MHETase is stable to pH, temperature, and for at least 12 days at room temperature.

**Conclusions:** We demonstrate the feasibility of using *S. cerevisiae* as a platform for the expression and surface display of PET degrading enzymes and predict that the whole-cell catalyst will viable alternatives to protein purification-based approaches for plastic degradation.

## Background

Since its invention over 70 years ago, plastic has become a major material for a wide range of items ranging from electronics components to clothing and packaging. It is currently estimated that over 360 million metric tonnes of plastics are produced every year [1,2]. In particular, the ease of production, cheap cost, and material versatility has made polyethylene terephthalate (PET) one of the most abundant plastics globally, with over 56 million metric tonnes produced every year, mainly for use in food packaging and textile fibers [1]. PET is easily produced by esterification of the petrochemicals ethylene glycol and terephthalic acid leading to the formation of polymers which can be easily molded into shape via melting processing, a process invented in the 1970’s [3].

Despite the enormous production of PET plastic, current solutions for waste management are lacking and it is estimated that at least 70% of total plastic is found as waste [1]. Two limitations account for the lack of effective plastic recycling solutions. First, recycling technologies for PET via physical or chemical processes leads to loss of material cohesion. Second, the current physical- and/or chemical-based methods of plastic recycling are not energy efficient as they often involve high temperatures and high pressures and often lead to the formation of hazardous by products, making them incompatible with environmentally conscious recycling approaches [1]. In addition, an increasing number of studies have shed light on the impact of plastic waste on animal and human health. Micro- and nano-plastics accumulate in animals from mollusc species to humans [4–6]. Although the physiological effects of these particles remain to be fully understood, recent studies suggest negative effects on biological functions such as oyster reproduction and hepatic lipid metabolism in mice [7,8]. Therefore, new methods for plastic waste management, remediation, and recycling are urgently needed.

Recently, enzymes capable of degrading PET plastic have been identified and engineered. In particular, the enzymes PETase and MHETase from the bacteria *Ideonella sakaiensis*, isolated from PET-polluted environmental samples, depolymerize PET into the two building blocks used for its synthesis, ethylene glycol (EG) and terephthalic acid (TPA) [9,10]. Importantly, EG and TPA obtained via enzymatic hydrolysis can be re-used for PET synthesis allowing complete and sustainable PET recycling [11,12].

Much current work has focused on improving PETase through protein engineering. Computational redesign of PETase has led to the development of thermostable variants of this mesophilic enzyme that are active at temperature close to the glass transition of PET, which increases polymer chain mobility to promote access to the ester linkages by the enzyme [11,13,14]. One recent and notable example of such approaches led to the identification of a new variant of PETase, dubbed FAST-PETase, containing 4 thermo-stabilizing mutations, boosting degradation efficiency up to 30-fold, and allowing degradation of entire post-consumer plastic containers in a matter of days [11]. Other studies have focused on identifying other PET degrading enzymes. Most examples involve enzymes from the cutinase, esterase and lipase families and were identified in bacteria and fungi. TfH (lipase), LCC, PHL7, HiC and Thc_Cut2 (cutinases) are among the other most promising PET-degrading enzymes and have been extensively characterized and engineered [12,15–19]. Although most of the research efforts have been focused on enzyme identification and enzyme engineering for use in the context of industrial processes using purified enzyme, microbe engineering for PET degradation and remediation has also been conducted. Heterologous expression of PET-degrading enzymes has been achieved in bacteria, yeast, and microalgae [20]. *Pseudomonas putida* has been extensively studied for PET degradation due to its ability to use EG as carbon source as well as for upcycling of TPA into higher value chemicals such as biodegradable plastics [21,22]. Other examples of TPA upcycling include conversion into catechol, muconic acid, glycolic acid, and vanillic acid [23,24]. More recently, *Pichia pastoris* was shown to be a suitable platform for expression of PETase and *Yarrowia lypolitica* was shown to naturally degrade PET and metabolize EG and TPA [25–28].

Despite the focus on PETase, MHETase is also a critical component of the enzymatic PET degradation process and is essential for converting the monohydroxyethyl terephthalate (MHET) product of the PETase reaction into TPA and EG. The PETase reaction products consist mainly of MHET, with TPA produced in small quantities if PETase is expressed alone [9]. MHET accumulation inhibits PET-hydrolysing enzymes [29,30] reducing their effectiveness, whereas dual systems such as fusion of PETase and MHETase improve PET hydrolysis [31]. Consequently, biological systems for MHETase expression and engineering are needed.

In this study, we establish a system to express MHETase from *Ideonella sakaiensis* on the surface of the yeast *Saccharomyces cerevisiae*. The resulting whole-cell biocatalyst allows conversion of MHET generated by PETase into TPA and EG (Figure 1A). We surveyed six potential surface display partners to identify a system that expresses MHETase at high density on the cell surface, and demonstrated that the resulting whole-cell catalyst hydrolyses a MHET analog without the need for purification of the MHETase enzyme. The activity of the MHETase whole-cell catalyst is similar to purified recombinant MHETase and is stable to alkaline pH, temperature, and for at least 12 days, a clear advantage over the purified enzyme. We anticipate that large-scale fermentation of the MHETase whole-cell biocatalyst will provide a low-cost source of MHETase suitable for PET plastic recycling, up-cycling, and remediation.

**Figure 1.**
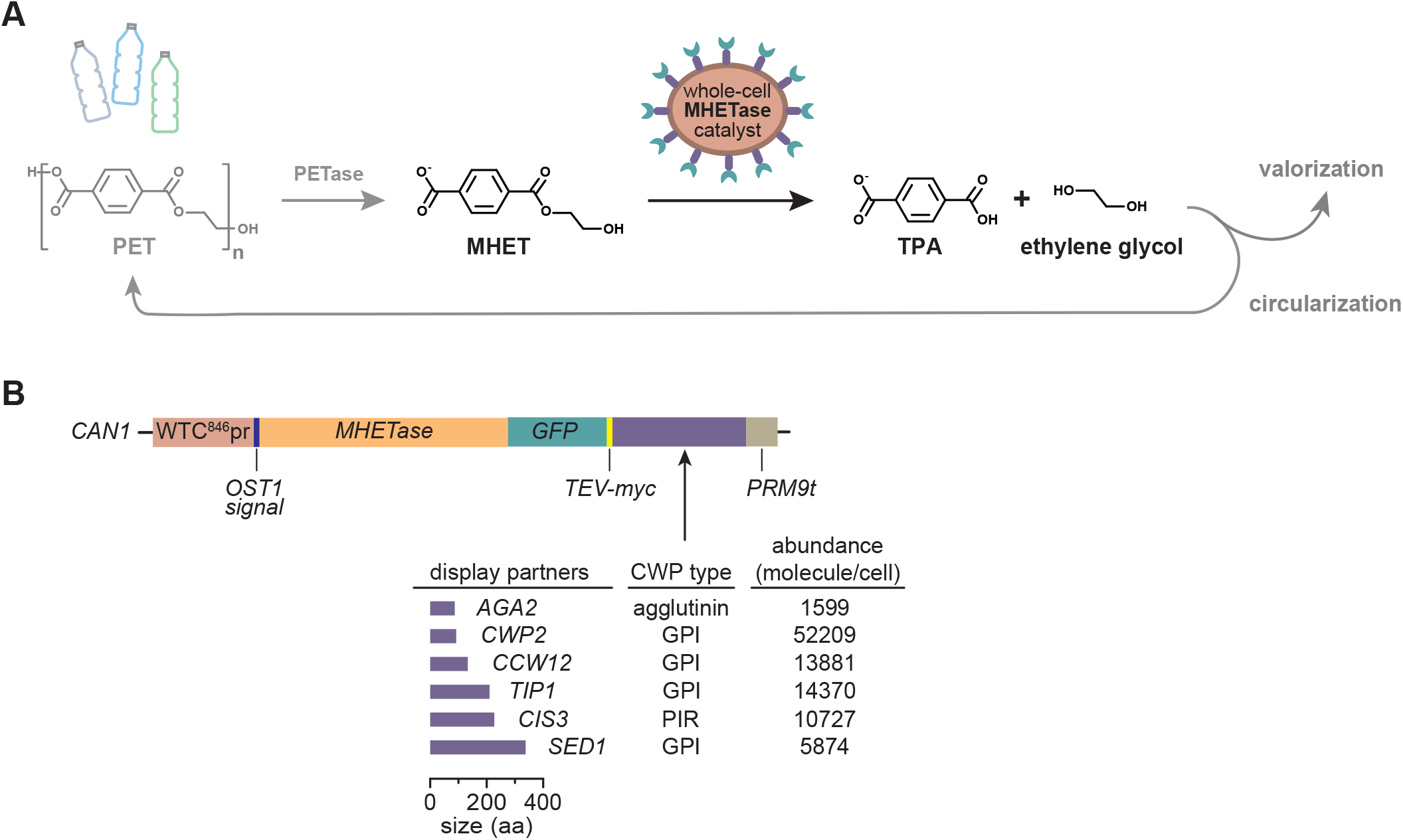
The MHETase whole-cell catalyst concept. **A**. The MHETase whole-cell catalyst performs the second step of the PET biodegradation pathway. In the first step, repeating units of MHET in the PET polymer are released by the enzyme PETase. MHET is then processed into TPA and ethylene glycol by the MHETase whole-cell catalyst. TPA and ethylene glycol can be used to synthesize new, virgin PET, bio-converted to high-value compounds or simply converted into biomass. **B**. Chimera design for surface display of MHETase. The coloured blocks represent the different components assembled to express MHETase (orange block) at the cell surface. Different cell wall proteins (CWPs; purple block) were fused to MHETase to identify the best design. Amino acid length is indicated, as is CWP type, and expression level for the different CWPs under their native promoters. Control chimeras lacking MHETase were also generated.

## Results and Discussion

### MHETase cell surface display modules

Our goal was to develop a system expressing the MHETase enzyme, from *I. sakaiensis*, in *S. cerevisiae* to process the product of PET-hydrolysis intermediate MHET (Figure 1A). Surface display is an ideal context for reactions with large substrates, like PET, that cannot translocate to the cell interior [32]. Additionally, surface display circumvents enzyme purification as a prerequisite for catalysis, avoids product contamination [33], facilitates reuse of the catalyst, and can increase catalyst stability [25,34]. We engineered a MHETase cell surface display system to probe these potential advantages relative to conventional enzyme expression and purification. The MHETase surface display system consists of an engineered transcription unit stably integrated at the *CAN1* locus driven by a doxycycline-inducible promoter (WTC846pr) to express MHETase fusion proteins (Figure 1B) [35]. The MHETase fusion contains *(i)* a secretion signal (from the *OST1* gene) fused to the N-terminus of the MHETase coding sequence, *(ii)* a yeast codon-optimized sequence of MHETase from *I. sakaiensis* followed by *(iii)* the coding sequence of GFP and *(iv)* the coding sequence of one of 6 display partners, namely *AGA2, CCW12, CIS3, CWP2, SED1* and *TIP1*, which encode yeast cell wall proteins, to allow anchoring of the MHETase protein chimera on the yeast surface (Figure 1B) [36]. The cell wall proteins used for anchoring at the cell surface were chosen to span different modes of covalent linkage to the cell wall, different molecular weights, and different expression levels (Figure 1B). We also designed modules driving secretion of soluble MHETase or intracellular MHETase, as controls.

### Efficient expression of MHETase display chimeras in vivo

Having successfully assembled the 8 MHETase modules, we measured protein expression using the GFP present in each chimeric protein. To accurately convert GFP fluorescence *in vivo* to protein abundance, we assembled a calibrating set of strains expressing GFP-tagged proteins with abundance ranging from 2.3×10^3^ to 7.5×10^5^ molecules/cell (Figure 2A) [37]. The correlation between protein abundance and normalized GFP intensity was excellent (R^2^ = 0.874, Figure 2A). Using the normalized GFP intensity measurements for the MHETase chimeras after 4 hours of induction, we calculated MHETase abundance in molecules/cell using the calibration curve (Figure 2B). MHETase chimeras were expressed at similar levels, ranging from 9.3 × 10^4^ (MHETase-Tip1) to 1.5 × 10^5^ (MHETase-Cis3) molecules/cell, corresponding to MHETase concentrations of 16 to 25 nM for cultures containing 10^8^ cells/mL (Figure 2B). The intracellular and the secreted MHETase were expressed at slightly higher levels (30 and 27 nM, respectively) compared to the MHETase display chimeras. When we assessed the expression level of the chimeras lacking MHETase, it became apparent that the MHETase sequences reduced protein expression, except for the Ccw12 fusion (Figure 2B). It is possible that the display partners, except Ccw12, do not tolerate additional cargo without some reduction of expression. Alternatively, there could be toxicity associated with MHETase expression. We compared growth of the strains expressing MHETase display chimeras with the growth of strains expressing GFP display chimeras. Only MHETase-Aga2 and MHETase-Cis3 resulted in a statistically-supported decrease in growth rate (Figure 2C), and the effect size was very small (approximately 5% decrease in growth rate). We conclude that MHETase expression is not toxic to the yeast platform.

**Figure 2.**
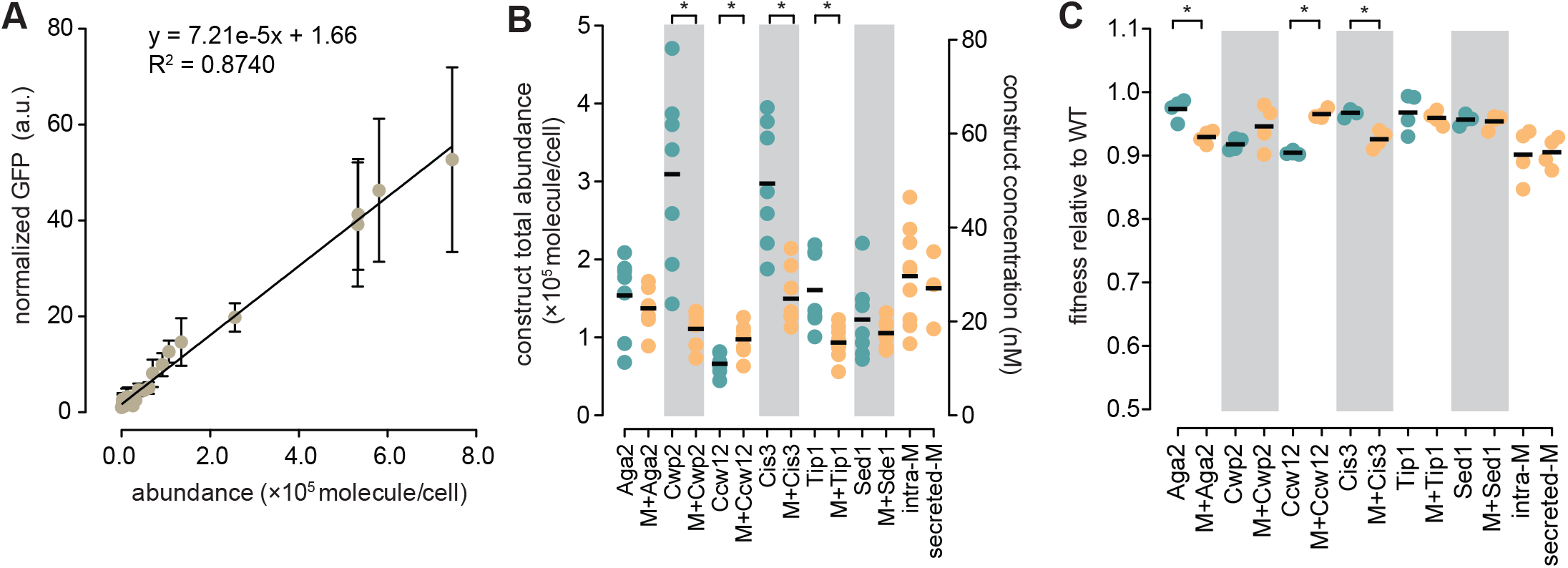
MHETase display constructs are efficiently expressed at minimal fitness cost. **A**. GFP calibration standards for measuring abundance of MHETase chimeras in molecules per cell. GFP-fusion strains spanning the range of molecules per cells were selected and GFP fluorescence was measured. The regression analysis line and equation are indicated. Bars indicate standard deviation; n ≥ 7. **B**. Abundance of the indicated surface display chimeras with (orange) or without (green) MHETase. Abundance was determined using GFP fluorescence after induction with doxycycline for 4 hours and converted to molecules per cell using the equation in A. Theoretical MHETase molarity was inferred from the molecule/cell data for a cell density of 10^8^ cells/ml (right y-axis). Horizontal bars indicate the means of the replicates. Asterisks indicate p-values ≤ 0.05 (unpaired Student’s t-test; n = 7). Intracellular MHETase (intra-M) and secreted MHETase (secreted-M) are indicated. **C**. Fitness of cells expressing the surface display chimeras. Cells expressing the indicated chimeras were grown in presence of doxycycline in YPD medium for 24h. Fitness is expressed as a ratio of the growth rate of each strain to that of the wild-type. Horizontal bars indicate the means of the replicates. Asterisks indicate p-values ≤ 0.05 (unpaired Student’s t-test; n = 4).

### An image analysis pipeline to quantify surface-displayed proteins

Total MHETase abundance does not accurately reflect the enzyme concentration at the cell surface. Secreted proteins can be retained intracellularly, reducing the amount of catalyst that is able to contact substrate outside of the cell, although display of cargos often has efficiency above 50% [38]. We developed a computational pipeline to analyse fluorescence microscopic images of yeast cells expressing surface display proteins to quantify the amount of protein at the cell surface relative to total protein expression. We imaged cells labelled with concanavalin A conjugated to Alexa Fluor 594 (conA-A594) which binds to glycoproteins in the cell wall. Cells were identified based on the conA-A594 fluorescence signal and concentric rings of 1 pixel width inside and outside the conA-A594-defined cell borders were segmented (Figure 3A). Fluorescence intensity was measured for each of the concentric rings. As shown in Figure 3B, the conA-A594 fluorescence signal followed a normal distribution between 0 and -9 pixels and peaked at -4 pixels, consistent with most of the signal being at the periphery of the cell and demonstrating that most of the cell wall signal is between 0 and -9 pixels inside the segmented cell object (Figure 3B). We repeated the analysis with conA-A594 labelled cells expressing Mrh1-GFP, a plasma membrane protein displaying a homogenous fluorescence signal at the cell periphery, as well as the MHETase intracellular chimera, and two additional intracellular GFP-tagged proteins, Tif2 and Rrp1A (Figure 3D). Tif2 and Rrp1A are expressed at 9.2 × 10^4^ and 1.4 × 10^5^ molecules/cell, respectively, similar to the expression levels of the MHETase display chimeras. As shown in Figure 3B, the fluorescence intensity profile for Mrh1-GFP closely followed that of conA-A594 consistent with Mrh1 residing at the cell periphery. Interestingly, the GFP signal for Mrh1-GFP peaked at the -5 pixels coordinate, while the conA-A595 signal peaked at the -4 pixels coordinate, indicating that our method can distinguish proteins at the plasma membrane from those at the cell wall. The Mrh1 C-terminus (including the GFP tag) is predicted to reside on the inner side of the plasma membrane (Figure S1) consistent with the GFP signal being more internal to the cell compared to the conA-A594 signal. By contrast, the fluorescence profile for the intracellular GFP-proteins did not resemble that of conA-A594 or Mrh1-GFP (Figure 3B). Instead, the fluorescence progressively increased from the -3 pixels ring and plateaued at -6 pixels, demonstrating that most of the signal is more internal as compared to the peak of fluorescence of both the plasma membrane and cell surface (Figure 3B). Even though the fluorescent signal was consistent with intracellular proteins, a significant amount of fluorescence signal was still present within the 0 to -9 ring, indicating bleed-through of intracellular fluorescence into the cell wall ring. For example, approximately 90% and 50% of the intracellular fluorescence intensity is still detected at the -4 and -5 rings, respectively, for all intracellular proteins (Figure 3C). Because the peak of cell surface fluorescence spanned the 0 to -9 pixel rings, we used this entire area to measure displayed abundance and corrected for intracellular fluorescence bleed-through (see Methods).

**Figure 3.**
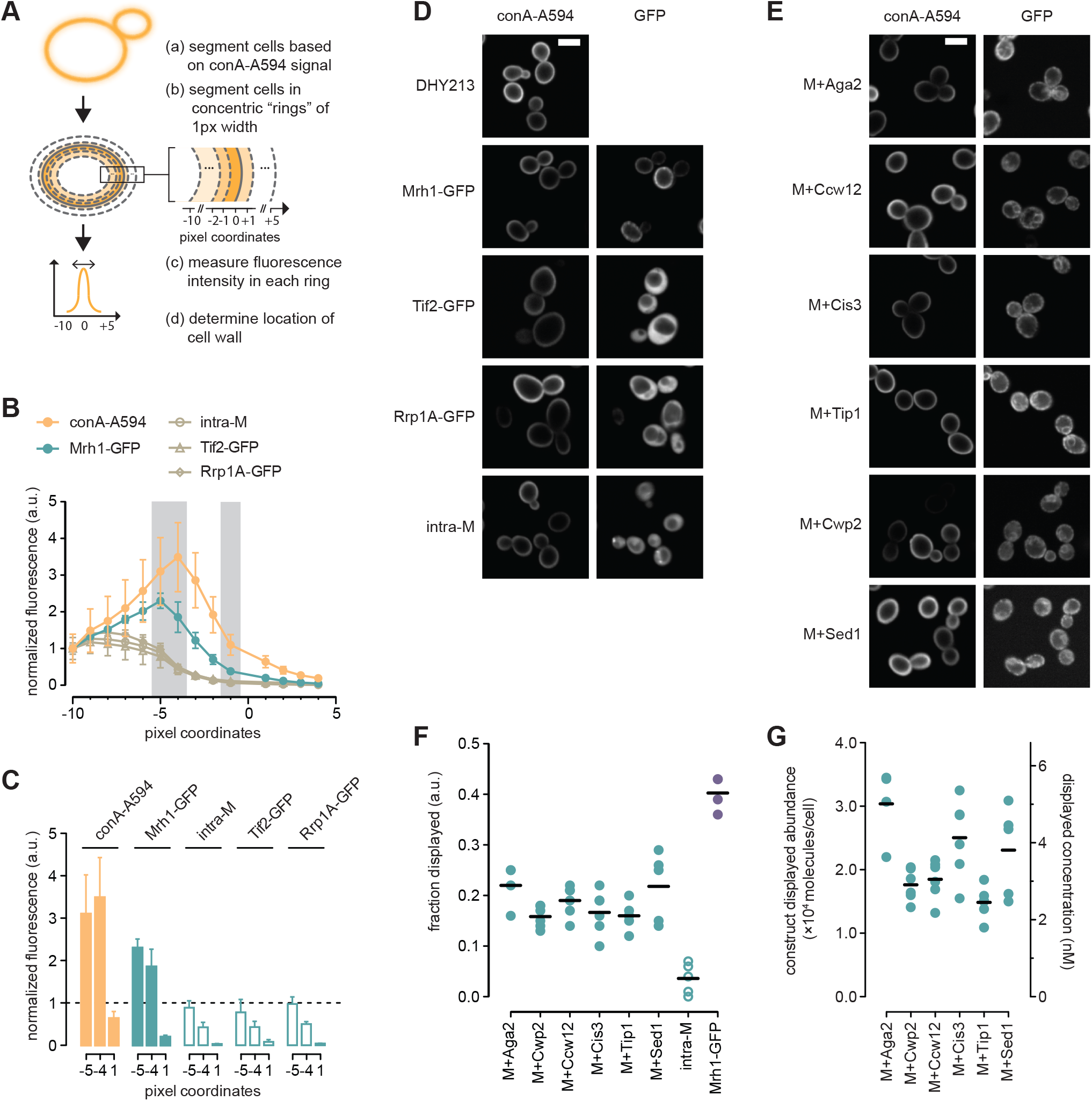
A microscopy-based method to measure MHETase cell surface display efficiency. **A**. Outline of the microscopy-based method to quantify GFP signal on the cell surface (see text for details). **B**. Mean fluorescence intensities at each pixel coordinate for the indicated strains. Bars indicate standard deviation. Analysis was performed on at least 40 cells in each replicate. conA-A594: n= 98, Mrh1-GFP and intra-cellular MHETase (intra-M): n = 4, Tif2-GFP and Rrp1A-GFP: n = 2. **C**. Comparison of mean fluorescence intensities for the -5, -4, and +1 pixel coordinates for the indicated strains (grey shading in B). Bars indicate standard deviation. **D, E**. Representative fluorescence micrographs for the strains in B and C and for the MHETase surface display chimeras. Scale bar: 5 μm. **F**. Fraction of MHETase chimeras displayed at the cell surface. Cells were induced for 4 hours, labelled with conA-A594 and imaged. The fraction of displayed chimera is plotted. Horizontal bars indicate the means of the replicates (n = 6). Each replicate included at least 200 cells. **G**. Abundance of the MHETase chimeras at the cell surface. The fraction of chimera displayed from panel F was used to calculate the cell surface abundance in molecules per cell. Theoretical construct molarity is indicated for a cell density of 10^8^ cells/ml. Horizontal bars indicate the means of the replicates (n = 6).

### MHETase is displayed efficiently at the cell surface

We determined the fraction of MHETase displayed at the cell surface by measuring the GFP signal at the cell surface relative to total GFP signal by analysis of fluorescence micrographs of cells expressing MHETase chimeras (Figure 3E). GFP intensity was integrated for the 0 to -9 pixel region and corrected for background and intracellular fluorescence bleed-through in the cell wall region and expressed as a ratio to total cell integrated GFP intensity. The analysis was performed on at least 200 individual cells in 6 replicates. As shown in Figure 3F, between 0.16 and 0.22 of the total MHETase was displayed at the cell surface, depending on the display partner. Next, using total abundance and displayed fraction data (Figure 2B, 3F), we calculated the displayed MHETase abundance in molecules/cell and in nanomolar concentration of enzyme for a suspension of cells at 10^8^ cells/ml. MHETase protein abundance ranged from 1.5 × 10^4^ (MHETase-Tip1) to 3.0 × 10^4^ (MHETase-Aga2) molecules/cell at the cell surface, corresponding to enzyme concentrations of 2.4 to 4.8 nM for 10^8^ cell/ml suspensions (Figure 3G). The MHETase-Aga2 and MHETase-Sed1 chimeras had the highest displayed fraction. The displayed protein abundance was more variable for MHETase-Aga2, MHETase-Sed1, and MHETase-Cis3 as compared to the other constructs, suggesting that cells might not display these chimeras uniformly. Although the displayed abundance for the MHETase-Aga2 (1.5 × 10^4^ molecules/cell) was consistent with those described for Aga1-Aga2 yeast surface display systems [32], none of the display partners moved more than 22% of total MHETase to the cell surface. Display efficiencies of over 50% have been described [38], and so we infer that there remains substantial room to improve the efficiency of our MHETase yeast surface display systems.

### Kinetic analysis of MHETase whole-cell catalysts

Having established that the MHETase constructs were expressed and displayed on the cell surface, we tested whether the MHETase whole-cell biocatalyst had the expected catalytic activity. MHETase activity is readily assayed with the colorimetric substrate MpNPT, and MHETase hydrolysis of MpNPT accurately reflects hydrolysis of MHET [29]. After 4 hours of induction, cells expressing MHETase chimeras were incubated with increasing concentrations of MpNPT and pNP formation was quantified. Enzymatic activity was normalized to 10^8^ cell/ml so that the different surface display chimeras could be compared. As shown in Figure 4A-F, all MHETase chimeras followed Michaelis-Menten kinetics. Differences in reaction rates and in substrate affinity were readily observable between chimeras, with MHETase-Aga2 performing poorly and MHETase-Tip1 having the highest reaction rate (Figure 4A-F). Importantly, cells expressing intracellular MHETase did not hydrolyse MpNPT, demonstrating that MpNPT is hydrolysed by the surface-displayed MHETase (Figure 4G). Recombinant MHETase produced in *E. coli* or secreted by yeast behaved similarly to the displayed MHETase chimeras (Figure 4H-I). Assays of 7 independent isolates of the MHETase-Tip1 chimera showed a high degree of reproducibility (Figure 4J), indicating that the whole-cell catalyst system is stable and robust to variation.

**Figure 4.**
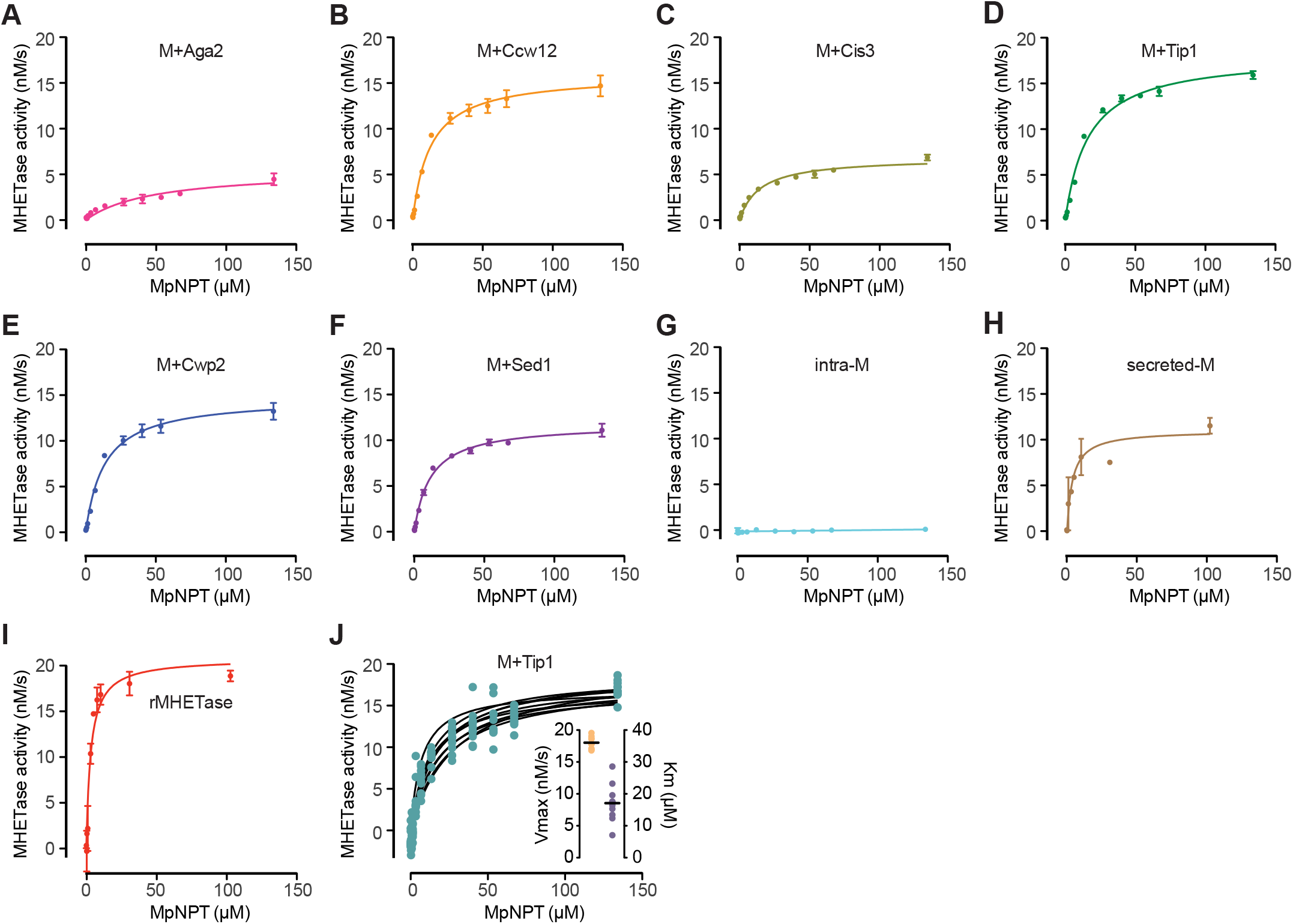
The MHETase whole-cell catalyst follows Michaelis-Menten kinetics. A through I. Michaelis-Menten plots for the MHETase chimeras and recombinant MHETase. For the displayed MHETase chimeras (A-G), cells were induced for 4 hours in YPD, rinsed twice and resuspended in 100mM phosphate buffer pH 7.5 prior to assaying MHETase activity by incubating with MpNPT at the indicated concentrations for 10 minutes at 24°C, followed by measuring absorbance at 405 nm. To allow comparison between samples of different cell density, MHETase activity was normalized to a cell density of 10^8^ cells/ml. For the recombinant and secreted enzyme (H-I), assays were performed under the same buffer and temperature conditions in the presence of the indicated MpNPT concentrations. Michaelis-Menten curves were fitted to the data. **J**. To test for system robustness, seven biological replicates of the MHETase-Tip1 fusion were assayed in parallel. Michaelis-Menten curves were fitted to each independent replicate (black lines). Vmax and Km were calculated from the fitted curves (inset).

To accurately compare the different MHETase chimeras to purified MHETase, kinetic parameters were calculated using the Michaelis-Menten plots, the enzyme concentration determined from total abundance, the display efficiency, and the cell culture density (Table 1). Again, differences between MHETase chimeras were readily observable. We found that the turnover number (k_cat_) for whole-cell catalysts were similar to MHETase purified from *E. coli* or MHETase secreted from yeast cells. MHETase-Tip1 k_cat_ was 68% of purified MHETase and 96% of secreted MHETase (Table 1). K_m_ values for the displayed chimeras were 3.6- to 15.7-fold greater than recombinant or secreted MHETase, indicating that surface display reduced the substrate affinity of MHETase. Consequently, catalytic efficiency for the whole-cell MHETase catalysts was also lower compared to recombinant or secreted MHETase. Lower substrate affinity and catalytic efficiency could be due to ectopic glycosylations that are typical of proteins transiting through the yeast secretory pathway [39]. However, the K_m_ of MHETase secreted from yeast was indistinguishable from that of purified MHETase, suggesting that glycosylation is not causing lower substrate affinity. We suggest that the reduced K_m_ of the surface displayed chimeras could reflect the environment of the yeast cell surface. As such, mutations that alter cell surface properties would be reasonable targets for improving the MHETase display platform. Interestingly, no correlation was evident between the activity of the different chimeras and expression at the cell surface, suggesting that the identity and the mode of cell surface anchoring itself might be responsible for the catalytic efficiency differences that we observe. Nevertheless, the displayed MHETase chimeras differ only modestly from purified MHETase, and our analyses highlight the importance of testing multiple surface display partners to identify chimeras with optimal catalytic properties.

**Table 1.** Enzymatic parameters for each of the display chimeras,secreted MHETase (secreted-M) and recombinant MHETase purified from *E. coli* (rMHETase).

|  | rMHETase | M+Aga2 | M+Ccw12 | M+Cis3 | M+Tip1 | M+Cwp2 | M+Sed1 | secreted-M |
| --- | --- | --- | --- | --- | --- | --- | --- | --- |
| **V<sub>max</sub>** (nM/s)<sup>a</sup> | 20.51 | 5.67 | 16.46 | 7.03 | 18.03 | 13.64 | 12.19 | 11.19 |
| **K<sub>m</sub>** (μM)<sup>a</sup> | 3.11 | 48.79 | 12.60 | 14.58 | 17.10 | 11.18 | 11.62 | 3.77 |
| **total [E]** (nM) | 1.90 | 22.78 | 16.22 | 24.83 | 15.50 | 18.43 | 17.52 | 1.47 |
| **display efficiency** | n.a. | 0.10 | 0.13 | 0.12 | 0.13 | 0.13 | 0.14 | n.a. |
| **displayed [E]** (nM) | n.a. | 2.35 | 2.18 | 3.04 | 2.07 | 2.49 | 2.50 | n.a. |
| **k<sub>cat</sub>** (s<sup>-1</sup>) | 10.79 | 2.41 | 7.55 | 2.31 | 8.72 | 5.49 | 4.87 | 7.61 |
| **efficiency** (μM<sup>-1</sup> s<sup>-1</sup>) | 3.47 | 0.05 | 0.60 | 0.16 | 0.51 | 0.49 | 0.42 | 2.02 |

<sup>a</sup> V<sub>max</sub> and K<sub>m</sub> were calculated from the Michaelis-Menton curves in Figure 4.

### The MHETase whole-cell catalyst is stable to alkaline pH, temperature and, time

We next established optimal reaction parameters for temperature and pH for the whole-cell catalyst. As shown in Figure 5A, enzymatic activity was optimal for all the chimeras at pH 7.5. At higher pH (pH 9.5 and 10.5), the system remained active, but activity was reduced by approximately 40 to 50%, which contrasts with purified MHETase which remained active at higher pH [29]. The differences observed for activity at pH 7.5 between the different chimeras (Figure 4) remained consistent across the pH range, with MHETase-Tip1 being the most active and MHETase-Aga2 displaying the lowest activity. Similarly, we assessed the effect of temperature on enzyme activity. As shown in Figure 6B, activity steadily increased and peaked at 45°C for all the chimeras. At 55°C, MHETase activity was lower. Therefore, of the tested temperatures, 45°C was optimal, with MHETase activity approximately 3-fold higher than at 24°C. Again, differences between chimeras were consistent across temperatures. Purified recombinant MHETase also showed optimal activity at 45°C, in agreement with previous characterizations of purified MHETase [29].

**Figure 5.**
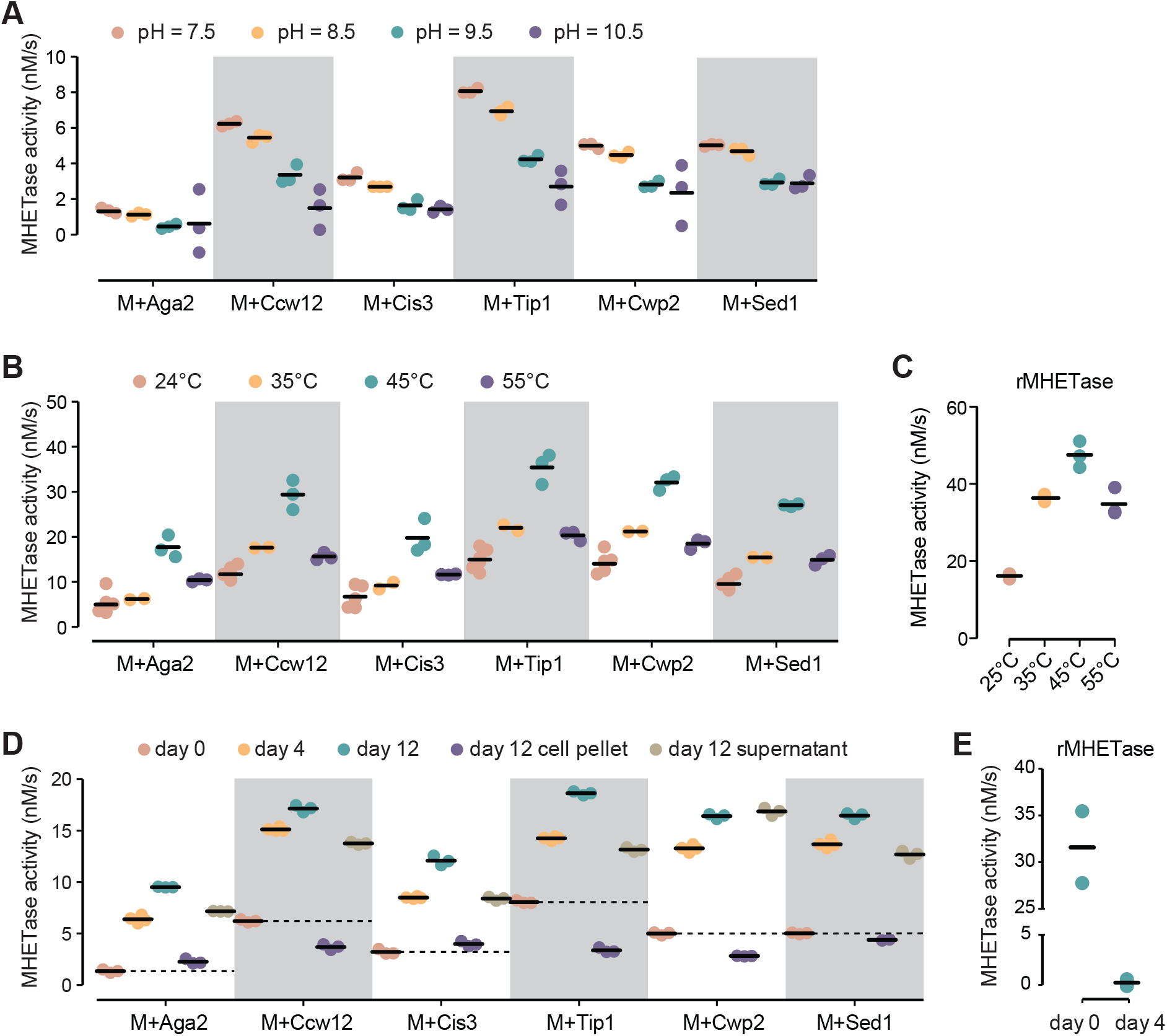
The MHETase whole-cell catalyst is stable to pH, temperature and time. **A**. MHETase activity at the indicated pH is plotted. MHETase activity was assayed with 26.8 μM MpNPT for 10 minutes at 24°C, followed by measuring absorbance at 405 nm. Activity was normalized to a cell density of 10^8^ cell/ml. Horizontal bars indicate the means of the replicates (n = 3). **B**. MHETase activity at the indicated temperatures is plotted. n = 3. **C**. Activity of purified MHETase at different temperatures is plotted. Purified enzyme was diluted to 2 nM and and assayed with 50 μM MpNPT. n = 3. **D**. MHETase activity of the whole-cell catalysts was assayed at day 0, 4, and 12 during incubation at room temperature. At day 12, the cell suspension, cell pellet, and supernatants were assayed. MHETase activity was normalized to a cell density of 10^8^ cell/ml at day 0. n = 3. **E**. Activity of purified MHETase over time. Purified enzyme was diluted to 2 nM and held at room temperature for 4 days. MHETase activity at day 4 was measured with 50 μM MpNPT at 25°C alongside a fresh aliquot of purified MHETase (day 0). n =2.

Surface display systems for PETase show little loss of enzyme activity over 7 days [25,40], whereas soluble PETase loses activity more rapidly. We compared the activity of surface displayed MHETase to soluble purified MHETase after 12 days at room temperature in phosphate buffer. Surprisingly, we observed that activity increased over time including for the cells expressing intracellular MHETase (Figure 5D, compare day 0 with day 4 or day 12). Interestingly, we noticed that cell count decreased over the same period of time by an average of 2- and 6.7-fold at day 4 and day 12, respectively, suggesting that cell lysis was occurring and that release of intracellular MHETase could be the mechanism by which activity is increasing. To test this hypothesis, we repeated enzymatic assay on precipitated cells washed with fresh phosphate buffer and on the supernatant of unwashed cells (Figure 5D). We observed strong activity in the supernatant, representing approximately 70-80% of enzymatic activity of the unwashed cell suspensions. We suggest that some caution is warranted in interpreting display stability results unless the whole-cell catalyst is washed prior to assay. Despite the finding that most of the MHETase activity at 12 days is no longer associated with the yeast cells, MHETase activity of the Aga2, Cis3, and Sed1 chimeras at the cell surface remained stable for 12 days (Figure 6D). By contrast, soluble purified MHETase was inactive after 4 days at room temperature demonstrating that the whole-cell catalyst is considerably more stable to prolonged incubation than the purified enzyme (Figure 6E).

## Conclusions

We have established a new system for degrading MHET, an important by-product of PET plastic degradation. Using a yeast surface display strategy and testing multiple display fusion partners, we demonstrate the production of MHETase at nanomolar concentrations in cell suspensions of moderate density (10^8^ cells/ml). We found that Aga2 was a poor display partner for MHETase. Although we note that display efficiency of MHETase-Aga2 was good, the K_m_ of MHETase-Aga2 was 4.4-fold higher than that of MHETase-Cwp2, and 16-fold higher than purified MHETase. We present alternative display partners for MHETase, including Tip1, Cwp2, and Sed1, that have suitable kinetic and display properties. MHETase whole-cell catalysts were stable for at least 12 days and retained activity up to 45°C. Stability gains relative to purified soluble MHETase when combined with time and cost savings realized by avoiding enzyme purification indicate that yeast surface display is a viable route for MHETase production. Finally, the yeast platform is amenable to synthetic biology, -omics, genetic, and artificial evolution strategies to improve the characteristics of the MHETase whole-cell catalyst.

## Methods

### Yeast maintenance and growth conditions

Yeast strains were maintained at 30°C in standard rich (YPD; 20 g/L peptone, 20g/L dextrose, 10 g/L yeast extract) or synthetic medium containing all amino acids (SDall; 6.7 g/L yeast nitrogen base, 20 g/L glucose). For MHETase induction, yeast strains were grown to saturation overnight in YPD and diluted 6-fold in fresh YPD containing doxycycline at a final concentration of 10 μg/mL. Cells were then grown for 4 hours with agitation at 30°C. Typical cell concentrations after 4 hours of induction were ∼10^8^ cell/mL. For MHETase secretion, the same induction scheme was used but cells were pre-grown in fully-supplemented synthetic medium (SDall) and induced in SDall containing 10 μg/ml doxycycline.

### Yeast strain construction

Yeast transformation was performed using the standard lithium acetate procedure. For CRISPR/Cas9 transformations, yeast cells were transformed using the pUB1306 plasmid (A kind gift of Elçin Ünal, originally generated by Gavin Schlissel and Jasper Rine) containing one of the following guide RNAs (CAN1 gRNA: GATACGTTCTCTATGGAGGA; OST1-GFP gRNA: TCATCGGCAATGGTCAGTAA) and transformants were selected on synthetic medium lacking uracil. URA+ transformants were then on 5-FOA medium to select against cells carrying the CRISPR/Cas9 plasmid. Transformants were validated by PCR and GFP expression was confirmed microscopically.

All strains were constructed in DHY213 (a derivative of BY4741 with higher sporulation efficiency and improved mitochondrial function [41]) and are listed in Table S1. To allow doxycycline induction of the WTC846 promoter [35], DHY213 was first modified by integrating the linearized FRP2370 plasmid (Addgene #127576), which encodes a cassette expressing the Tet repressor, yielding strain RLKY218 (Table S1). All subsequent strains were constructed in the RLKY218 background via CRISPR/Cas9 mediated assembly of PCR fragments at the *CAN1* locus. A first set of strains with the following construct architecture was generated: *WTC846pr-OST1ss-GFP-display_partner-PRM9ter*. WTC846pr is a strong doxycycline inducible promoter, *OST1ss* is the Ost1 endoplasmic reticulum translocation signal to allow for efficient secretion [36], GFP is the yeast codon-optimized monomeric GFP [42,43], *display_partner* is the coding sequence of one of *SED1, AGA2, CCW12, CWP2, CIS3* or *TIP1* lacking their respective secretion signals, and *PRM9ter* is the terminator region of *PRM9* (Table S2). The display partner sequences were codon optimized to minimize chances of recombination between the endogenous loci and the synthetic constructs, which were integrated at *CAN1*. Codon optimization was performed using the “Optimize codon” function of Benchling (https://www.benchling.com/) using *Saccharomyces cerevisiae* as “Organism”. This first set of strains was then used as platform for integration of the yeast codon optimized MHETase gene from *I. sakaiensis* (devoid of its endogenous secretion signal) between the *OST1ss* and the *msGFP* sequence (Table S2). All DNA sequences described here are provided in the Table S2.

### Measuring MHETase total protein abundance

Expression was induced as described above. After 4h of induction, cells were washed twice with sterile water and resuspended in the same volume of sterile water. 200 μL of cells were transferred into a clear 96-well plate and GFP fluorescence intensity was measured. The same cell suspension was diluted 10 times and used to measure optical density at 600nm (OD_600_). All measurements were made using a CLARIOstar (BMG LABTECH) plate reader. For each strain, GFP intensity was first corrected for cell mass by dividing GFP intensity by OD_600_ (*GFP*_*corr*_). *GFP*_*corr*_ values were then expressed as a ratio (*GFP*_*norm*_) between *GFP*_*corr*_ for a given GFP expressing strain and *GFP*_*corr*_ obtained for a GFP negative control strain (DHY213).

To establish a GFP standard curve, the following strains were obtained from the GFP strain collection [37,44]: *PEX21-GFP, FMP23-GFP, MDL2-GFP, PER1-GFP, LPX1-GFP, YML007CA-GFP, RAI1-GFP, SPI1-GFP, RTG2-GFP, MOT2-GFP, RRP15-GFP, RET2-GFP, GCN20-GFP, RPC40-GFP, NEW1-GFP, ARB1-GFP, OLA1-GFP, RPL2A-GFP, PMP2-GFP, STM1-GFP, TIF2-GFP, HTB2-GFP, RPS1B-GFP, RPP1A-GFP, SSA2-GFP, SSA1-GFP, TEF2-GFP, TEF1-GFP, PDC1-GFP, TDH3-GFP* and their GFP fluorescence intensity was measured. Regression analysis was performed with *GFP*_*norm*_ values for the GFP strains and the median molecules/cell data from Ho *et al* [37,44], using GraphPad Prism 5. *GFP*_*norm*_ values obtained for the various surface display constructs were then used to calculate their respective abundances using the regression equation determined from the GFP standard curve.

### Measuring MHETase cell surface abundance

Cells were induced in YPD as described above. After 4 hours of induction cells were washed in sterile water twice and resuspended in water containing 10 μm/mL concanavalin A conjugated with Alexa Fluor 594 (Thermo Fisher Scientific) and incubated at room temperature for 1 hour. GFP and Alexa Fluor 594 imaging was performed on an Opera Phenix (Perkin Elmer) high-throughput confocal microscope at a focal height of 1.5 μm using 488 nm and 561 nm excitation lasers and 500-550 nm, 570-630 nm bandpass emission filters. Images were analyzed with CellProfiler 3.1.9 (https://cellprofiler.org/) using the custom pipeline provided in the supplementary material.

To determine the position of cell surface with respect to the outline of the segmented cell objects, cells were first identified and segmented using Alexa Fluor 594 fluorescence images. Cell objects were further segmented into 10 inward and 4 outward concentric rings of one pixel width except for the most inward ring which represented the remaining inner portion of the cell. Median fluorescence was determined in each ring and corrected for background fluorescence before being normalized by the signal of most inner portion of the cell. Cell wall signal was determined as the area of strongest concanavalin A signal, which spanned a ring of 9 pixels width inside the cell object (Figure 3B, conA-A594 curve). This analysis was also performed on cells expressing known intracellular GFP-tagged proteins (Rrp1a-GFP, Tif2-GFP and intra-M chimera) to determine the average fraction of inner fluorescence signal spreading into each of the cell wall rings defined above (Figure 3B). The fraction of inner fluorescence was termed *FB*_*i*_ (Fluorescence Bleed, where *i* represents a given 1-pixel width ring). This parameter was used in the analysis below.

To determine the abundance of MHETase at the cell surface, the GFP intensity was integrated for the entire cell object and for the 9 inner rings closest to the cell object outer edge and expressed as a ratio of integrated GFP in the cell wall ring over the integrated GFP for the entire cell. We refer to this ratio as the fraction of GFP displayed or display efficiency. To account for background fluorescence and intracellular bleed-through fluorescence, two normalizations were applied before calculating the fraction of GFP displayed. First, all raw integrated GFP values were corrected for background fluorescence as follows: *GFPint*_*corr*1_ = *GFPinti* − (*GFPmed*_*Backd*_ × *P*_*i*_), where *GFPint*_*i*_ is the raw GFP integrated value for a given ring or the total cell, *GFPmed*_*backd*_ is the median background fluorescence determined from an area of the image with no cells and *P*_*i*_ the number of pixels in the area considered (ring or total cell). Second, bleed-through fluorescence was also taken into account for integrated GFP values of each of the 9 cell wall rings, as follows: *GFPint*_*corr*2_ = *GFPint*_*corr*1_ − (*GFPmed*_*iinner*_ × *FB*_*i*_ × *P*_*i*_), where *GFBmed*_*inner*_ is the background corrected median GFP fluorescence intensity for the inner part of the cell, *FB*_*i*_ is the fluorescence bleed-through correction factor for the area considered, as determined above, and *P*_*i*_ the number of pixels in the ring area considered. Displayed ratio was then calculated as the sum of *GFPint*_*corr2*_ values from the cell wall rings and divided by *GFPint*_*corr1*_ obtained for the total cell. At least 200 cells were analyzed in each technical (n=2) and biological replicates (n=3).

### Measurement of strain fitness

Fitness was measured as previously described [45]. Briefly, cells were grown to saturation overnight and diluted 100-fold in 200 μL of fresh YPD with or without doxycycline (10 μg/mL) in a transparent 96-well plate. OD600 was monitored every 15 minutes in a Genios Tecan plate reader. Growth rate was determined in R (https://www.r-project.org/). Fitness was calculated as the ratio of the growth rate of the experimental strain to that of the parental strain (DHY213).

### MHETase activity measurement with the whole-cell biocatalyst

Induced cells were washed twice in sterile water and resuspended in same volume with 111 mM phosphate buffer at pH 7.5, 8.5, 9.5 or 10.5. Cell concentration was determined using a Beckman-Coulter Counter Z1 equipped with a 100 μm aperture tube using a particle lower threshold limit of 4 μm. 270 μL of cells were mixed with 30 μL of MpNPT (CAS #1137-99-1, Toronto Research Chemicals) at ten times the final concentration in DMSO, and reaction was allowed to proceed for 10 minutes. The reaction was stopped by separating the cells from the reaction with a 96-well filter plate (AcroPrep, Pall) mounted on a vacuum device (NucleoVac 96, Macherey-Nagel). Alternatively, miniprep columns were used for filtering (PuroSPIN MINI, Luna Nanotech). 120 μL of filtered reaction was then transferred into a clear 384-well plate, to increase the light pathlength, and *para*-nitrophenol (pNP) concentration was determined by measuring absorbance at 405nm in a CLARIOstar plate reader (BMG LABTECH). Each run included an MpNPT autohydrolysis control (MpNPT diluted in phosphate buffer only). The molar extinction coefficients at 407nm for pNP at the different pH’s were calculated from Biggs (1954) [46] and are provided in Figure S2. All reactions were performed at 24°C unless specified otherwise. To assess activity at different temperatures, cells were pre-incubated in a water bath at the given temperature for 10 minutes before addition of the substrate and held at the same temperature after addition of MpNPT. To test the stability of the whole-cell biocatalyst, induced cells were resuspended in phosphate buffer pH 7.5 and held for 12 days at room temperature without agitation.

### Purification, quantification, and activity measurement of recombinant MHETase from E. coli

Recombinant MHETase was purified as described previously [29] with some modifications. *E. coli* Shuffle T7 express cells were transformed with pCOLDII-MHETase vector [29] and selected on agar plates containing 100 μg/mL carbenicillin at 30°C. Single colonies were inoculated into liquid growth medium containing carbenicillin and protein expression was induced as follows. 1L cultures were grown to an OD of ∼0.5 at 30°C, then rapidly cooled in an ice bath to ∼10°C. Isopropyl β-D-1-thiogalactopyranoside (IPTG) was added to a final concentration of 1 mM, and cultures were incubated overnight at 16°C with shaking. Cell pellets were collected by centrifugation at 16,770 g at 4°C, resuspended in 50mM Tris-HCl (pH 7.5), 100 mM NaCl, 10 mM imidazole, 1 mM DTT, and protease inhibitors (2 μg/mL aprotonin, 10 μM bestatin, 10 μM leupeptin, 1 μM pepstatin, and 0.5 mM PMSF), and lysed by sonication then clarified by ultracentrifugation (4°C, 142,000 g, 1 hour). The clarified lysates were loaded onto a 5 mL His-Trap FF column (Cytiva), washed with 50 mM Tris-HCl (pH 7.5), 100 mM NaCl, 20 mM imidazole and 1 mM DTT, and then eluted in 50 mM Tris-HCl (pH 7.5), 100 mM NaCl and 500 mM imidazole. Peak fractions were pooled and diluted with 25 mM Tris-HCl (pH 7.5) to a final concentration of ∼50 mM NaCl before loading onto a 5 mL HiTrap Q HP column (Cytiva) pre-equilibrated in 25 mM Tris-HCl (pH 7.5) and 50 mM NaCl. The column was then washed using 10 column volumes of 25 mM Tris-HCl (pH 7.5) and 50 mM NaCl, followed by a 0.05-1 M NaCl salt gradient over 10 column volumes. As most of the recombinant MHETase eluted in the wash, the wash fraction was concentrated to a final volume of ∼500 μL with an Ultra-15 10kDa MWCO centrifugal concentrator (Amicon) and then loaded onto a Superdex 75 Increase 10/300 GL column (Cytiva). Recombinant MHETase was then eluted in 20 mM Tris-HCl (pH 7.5) and 150 mM NaCl at 0.5 mL/min and peak fractions were pooled. Protein purity was assessed by SDS-PAGE (Figure S3) and protein concentration was measured spectrophotometrically using ε_280_ = 102,330 M^-1^cm^-1^. Protein aliquots were snap-frozen prior to being stored at -80°C.

Recombinant MHETase activity was measured as described previously [29] in 100 mM sodium phosphate buffer (pH 7.5) at 24°C. Enzymatic parameters were similar to published data for MHETase using MpNPT as substrate [9,29]. To assess activity at different temperatures, MHETase in 100 mM phosphate buffer pH 7.5 was pre-incubated in a water bath at the given temperature for 20 minutes before addition of the substrate and held at the same temperature after addition of MpNPT. The enzyme was freshly thawed before each assay. To determine stability over time, the recombinant enzyme was kept at room temperature in 100 mM sodium phosphate buffer (pH 7.5) for 4 days without shaking.

### Purification, quantification, and activity measurement of MHETase secreted from yeast

Cultures of RLKY245 (intracellular MHETase control) and RLKY247 (*OST1-MHETase-GFP*) were grown overnight in SDall at 30°C. The overnight culture was then induced by the addition of 10 μg/ml doxycycline as described above. After 4 hours of induction, cells were centrifuged at 3500 rpm for 5 minutes at room temperature, and the supernatant was collected and kept on ice throughout the remainder of the procedure. The supernatant was concentrated to a final volume of ∼300 μL, and buffer exchanged to 100 mM sodium phosphate buffer pH 7.5 (Amicon Ultra-4, Millipore Sigma). The concentrated sample was stored at 4°C for a maximum of one week.

MHETase concentration was measured by ELISA. Samples were diluted 2-, 4- and 8-fold in sodium phosphate pH 7.5. Clear flat-bottom Immuno Nonsterile 96-well plates (Thermo Fisher Scientific) were coated with the samples, or with serial dilutions of purified GFP (Invitrogen; concentration range of 0.1-50 ng/mL) at 4°C overnight. The coating solution was then removed and 200 μL of blocking buffer (1x PBS, 3% non-fat milk, 0.1% Tween-20) was added to each well and incubated at room temperature for 1 hour. After removal of the blocking solution 100 μL of anti-GFP (Living Colors GFP monoclonal antibody, Clontech) diluted 1:10,000 in antibody solution (1x PBS, 1% non-fat milk, 0.1% Tween-20) was added to each well and incubated at room temperature for 2 hours. Plates were washed 3 times for 5 minutes each with PBS-T (1x PBS, 0.1% Tween 20). After removing the wash solution, 50 μL of anti-mouse-HRP (Pierce) diluted 1:10,000 in antibody solution was added to the plates, and incubated for 1 hour at room temperature. Plates were then washed 3 times for 5 minutes each with PBS-T at room temperature. After removing the wash solution, 100 μL of TMB substrate (Thermo Fisher Scientific) was added to each well. The reaction was incubated in the dark at room temperature for a maximum of 10 minutes and stopped by adding 50 μL of 2 N HCl to each well. Absorbance was measured at 450nm on a microplate reader (CLARIOstar, BMG LabTech) and measurements from RLKY245 supernatant were used as the negative control for the measurements of the RLKY247 supernatant. MHETase activity was assayed as described above for the recombinant MHETase purified from *E. coli*.

## Supporting information

CellProfiler pipeline

## Acknowledgements

The authors thank Gottfried Palm and Uwe Bornscheuer for the kind gift of pColdII-MHETase, Elçin Ünal for the kind gift of pUB1306, and Noor Hashem and Thomas Zheng for the original yeast PET degradation concept. We are grateful to work on the lands of the Mississaugas of the Credit, the Anishnaabeg, the Haudenosaunee and the Wendat peoples, land that is now home to many diverse First Nations, Inuit, and Métis peoples.

## Author contributions

Conceptualization, RLK, VS, and GWB; Methodology, RLK, VS, MWF, BH, BJP, and GWB; Protein purification, MWF and BJP; Experimentation, RLK, MWF, and BH; Strain engineering, VS, JB and SP; Formal Analysis, RLK, MWF, BH, and GWB; Funding acquisition, HDMW and GWB; Writing, RLK, MWF, BH, and GWB; Review & Editing, RLK, VS, MWF, BH, BJP, JB, SP, HDMW, and GWB.

## Availability of data and materials

All data supporting the conclusions of this study are included within the article and its additional files.

## Declarations

### Ethics approval and consent to participate

Not applicable

### Consent for publication

Not applicable

## References

1. Soong YHV, Sobkowicz MJ, Xie D. Recent Advances in Biological Recycling of Polyethylene Terephthalate (PET) Plastic Wastes. Bioengineering. 2022;9.

2. Plastics - the Facts 2021. http://www.plasticseurope.org/knowledge-hub/plastics-the-facts-2021/.

3. Wyeth NC, Al-E, Convers N, Ronald W, Roseveare N. Biaxially oriented poly(ethylene terephthalate) bottle. 1970.

4. Toussaint B, Raffael B, Angers-Loustau A, Gilliland D, Kestens V, Petrillo M, et al. Review of micro- and nanoplastic contamination in the food chain. Food Additives & Contaminants: Part A. 2019;36:639–73.

5. Ragusa A, Svelato A, Santacroce C, Catalano P, Notarstefano V, Carnevali O, et al. Plasticenta: First evidence of microplastics in human placenta. Environ Int. Environ Int. 2021;146.

6. Schwabl P, Koppel S, Konigshofer P, Bucsics T, Trauner M, Reiberger T, et al. Detection of Various Microplastics in Human Stool: A Prospective Case Series. Ann Intern Med. Ann Intern Med. 2019;171:453–7.

7. Sussarellu R, Suquet M, Thomas Y, Lambert C, Fabioux C, Pernet MEJ, et al. Oyster reproduction is affected by exposure to polystyrene microplastics. Proc Natl Acad Sci U S A. 2016;113:2430–5.

8. Lu L, Wan Z, Luo T, Fu Z, Jin Y. Polystyrene microplastics induce gut microbiota dysbiosis and hepatic lipid metabolism disorder in mice. Sci Total Environ. Sci Total Environ. 2018;631– 632:449–58.

9. Yoshida S, Hiraga K, Takehana T, Taniguchi I, Yamaji H, Maeda Y, et al. A bacterium that degrades and assimilates poly(ethylene terephthalate). Science. 2016;351:1196–9.

10. Joo S, Cho IJ, Seo H, Son HF, Sagong HY, Shin TJ, et al. Structural insight into molecular mechanism of poly(ethylene terephthalate) degradation. Nat Commun. Nat Commun. 2018;9.

11. Lu H, Diaz DJ, Czarnecki NJ, Zhu C, Kim W, Shroff R, et al. Machine learning-aided engineering of hydrolases for PET depolymerization. Nature. 2022;604:662–7.

12. Tournier V, Topham CM, Gilles A, David B, Folgoas C, Moya-Leclair E, et al. An engineered PET depolymerase to break down and recycle plastic bottles. Nature. 2020;580:216– 9.

13. Cui Y, Chen Y, Liu X, Dong S, Tian Y, Qiao Y, et al. Computational Redesign of a PETase for Plastic Biodegradation under Ambient Condition by the GRAPE Strategy. ACS Catal. 2021;11:1340–50.

14. Son HF, Cho IJ, Joo S, Seo H, Sagong HY, Choi SY, et al. Rational Protein Engineering of Thermo-Stable PETase from Ideonella sakaiensis for Highly Efficient PET Degradation. ACS Catal. 2019;9:3519–26.

15. Müller RJ, Schrader H, Profe J, Dresler K, Deckwer WD. Enzymatic degradation of poly(ethylene terephthalate): Rapid hydrolyse using a hydrolase from T. fusca. Macromol Rapid Commun. 2005;26:1400–5.

16. Sulaiman S, You DJ, Kanaya E, Koga Y, Kanaya S. Crystal structure and thermodynamic and kinetic stability of metagenome-derived LC-cutinase. Biochemistry. 2014;53:1858–69.

17. Sulaiman S, Yamato S, Kanaya E, Kim JJ, Koga Y, Takano K, et al. Isolation of a novel cutinase homolog with polyethylene terephthalate-degrading activity from leaf-branch compost by using a metagenomic approach. Appl Environ Microbiol. 2012;78:1556–62.

18. Herrero Acero E, Ribitsch D, Steinkellner G, Gruber K, Greimel K, Eiteljoerg I, et al. Enzymatic surface hydrolysis of PET: Effect of structural diversity on kinetic properties of cutinases from Thermobifida. Macromolecules. 2011;44:4632–40.

19. Sonnendecker C, Oeser J, Richter PK, Hille P, Zhao Z, Fischer C, et al. Low Carbon Footprint Recycling of Post-Consumer PET Plastic with a Metagenomic Polyester Hydrolase. ChemSusChem. 2021.

20. Dissanayake L, Jayakody LN. Engineering Microbes to Bio-Upcycle Polyethylene Terephthalate. Front Bioeng Biotechnol. 2021;9.

21. Kenny ST, Runic JN, Kaminsky W, Woods T, Babu RP, Keely CM, et al. Up-cycling of PET (polyethylene terephthalate) to the biodegradable plastic PHA (polyhydroxyalkanoate). Environ Sci Technol. 2008;42:7696–701.

22. Werner AZ, Clare R, Mand TD, Pardo I, Ramirez KJ, Haugen SJ, et al. Tandem chemical deconstruction and biological upcycling of poly(ethylene terephthalate) to β-ketoadipic acid by Pseudomonas putida KT2440. Metab Eng. 2021;67:250–61.

23. Sadler JC, Wallace S. Microbial synthesis of vanillin from waste poly(ethylene terephthalate). Green Chemistry. 2021;23:4665–72.

24. Kim HT, Kim JK, Cha HG, Kang MJ, Lee HS, Khang TU, et al. Biological Valorization of Poly(ethylene terephthalate) Monomers for Upcycling Waste PET. ACS Sustain Chem Eng. 2019;7:19396–406.

25. Chen Z, Wang Y, Cheng Y, Wang X, Tong S, Yang H, et al. Efficient biodegradation of highly crystallized polyethylene terephthalate through cell surface display of bacterial PETase. Science of The Total Environment. 2020;709:136138.

26. Gamerith C, Vastano M, Ghorbanpour SM, Zitzenbacher S, Ribitsch D, Zumstein MT, et al. Enzymatic degradation of aromatic and aliphatic polyesters by P. pastoris expressed cutinase 1 from Thermobifida cellulosilytica. Front Microbiol. 2017;8:938.

27. da Costa AM, de Oliveira Lopes VR, Vidal L, Nicaud JM, de Castro AM, Coelho MAZ. Poly(ethylene terephthalate) (PET) degradation by Yarrowia lipolytica: Investigations on cell growth, enzyme production and monomers consumption. Process Biochemistry. 2020;95:81–90.

28. Kosiorowska KE, Biniarz P, Dobrowolski A, Leluk K, Miroñczuk AM. Metabolic engineering of Yarrowia lipolytica for poly(ethylene terephthalate) degradation. Science of The Total Environment. 2022;831:154841.

29. Palm GJ, Reisky L, Böttcher D, Müller H, Michels EAP, Walczak MC, et al. Structure of the plastic-degrading Ideonella sakaiensis MHETase bound to a substrate. Nat Commun. 2019;10.

30. Barth M, Oeser T, Wei R, Then J, Schmidt J, Zimmermann W. Effect of hydrolysis products on the enzymatic degradation of polyethylene terephthalate nanoparticles by a polyester hydrolase from Thermobifida fusca. Biochem Eng J. 2015;93:222–8.

31. Knott BC, Erickson E, Allen MD, Gado JE, Graham R, Kearns FL, et al. Characterization and engineering of a two-enzyme system for plastics depolymerization. Proc Natl Acad Sci U S A. 2020;117:25476–85.

32. Andreu C, del Olmo M. Yeast arming systems: pros and cons of different protein anchors and other elements required for display. Appl Microbiol Biotechnol. 2018;102:2543–61.

33. Hartmann M, Kostrov X. Immobilization of enzymes on porous silicas-benefits and challenges. Chem Soc Rev. 2013;42:6277–89.

34. Yuzbasheva EY, Yuzbashev T v., Perkovskaya NI, Mostova EB, Vybornaya T v., Sukhozhenko A v., et al. Cell surface display of Yarrowia lipolytica lipase Lip2p using the cell wall protein YlPir1p, its characterization, and application as a whole-cell biocatalyst. Appl Biochem Biotechnol. 2015;175:3888–900.

35. Azizoglu A, Brent R, Rudolf F. A precisely adjustable, variation-suppressed eukaryotic transcriptional controller to enable genetic discovery. Barkai N, editor. Elife. 2021;10:e69549.

36. Fitzgerald I, Glick BS. Secretion of a foreign protein from budding yeasts is enhanced by cotranslational translocation and by suppression of vacuolar targeting. Microb Cell Fact. 2014;13:125.

37. Ho B, Baryshnikova A, Brown GW. Unification of Protein Abundance Datasets Yields a Quantitative Saccharomyces cerevisiae Proteome. Cell Syst. 2018;6:192-205.e3.

38. van der Vaart JM, te Biesebeke R, Chapman JW, Toschka HY, Klis FM, Verrips CT. Comparison of cell wall proteins of Saccharomyces cerevisiae as anchors for cell surface expression of heterologous proteins. Appl Environ Microbiol. 1997;63:615–20.

39. Delic M, Valli M, Graf AB, Pfeffer M, Mattanovich D, Gasser B. The secretory pathway: exploring yeast diversity. FEMS Microbiol Rev. 2013;37:872–914.

40. Jia Y, Samak NA, Hao X, Chen Z, Wen Q, Xing J. Hydrophobic cell surface display system of PETase as a sustainable biocatalyst for PET degradation. Front Microbiol. 2022;13:1005480.

41. Harvey CJB, Tang M, Schlecht U, Horecka J, Fischer CR, Lin H-C, et al. HEx: A heterologous expression platform for the discovery of fungal natural products. Sci Adv. 2018;4:eaar5459.

42. Kaishima M, Ishii J, Matsuno T, Fukuda N, Kondo A. Expression of varied GFPs in Saccharomyces cerevisiae: codon optimization yields stronger than expected expression and fluorescence intensity. 2016;6.

43. Cinelli RAG, Ferrari A, Pellegrini V, Tyagi M, Giacca M, Beltram F. The Enhanced Green Fluorescent Protein as a Tool for the Analysis of Protein Dynamics and Localization: Local Fluorescence Study at the Single-molecule Level. Photochem Photobiol. 2000;71:771–6.

44. Huh WK, Falvo JV, Gerke LC, Carroll AS, Howson RW, Weissman JS, et al. Global analysis of protein localization in budding yeast. Nature. 2003;425:686–91.

45. Loll-Krippleber R, Brown GW. P-body proteins regulate transcriptional rewiring to promote DNA replication stress resistance. Nat Commun. 2017;8.

46. Biggs AI. A spectrophotometric determination of the dissociation constants of p-nitrophenol and papaverine. Transactions of the Faraday Society. 1954;50:800–2.

