## Supplemental Figures for "Development of a yeast whole-cell biocatalyst for MHET conversion into terephthalic acid and ethylene glycol"

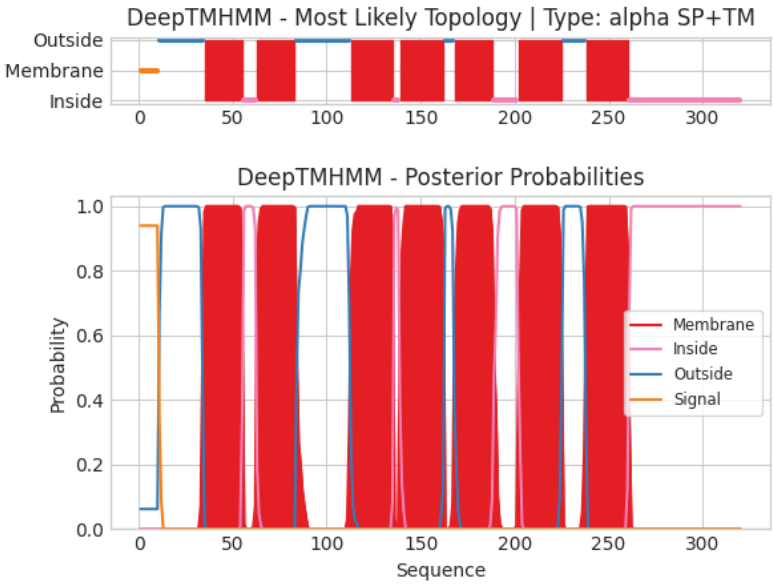


**Figure S1. Mrh1 transmembrane topology prediction.** Results obtained from the DeepTMHMM application (dtu.biolib.com/DeepTMHMM, accessed September 28, 2022) for the Mrh1 protein. Top panel: protein domain orientation relative to the inner and outer part of the cytoplasmic membrane. Bottom panel: probability associated with the inner and outer orientation for each protein domain.

**A**

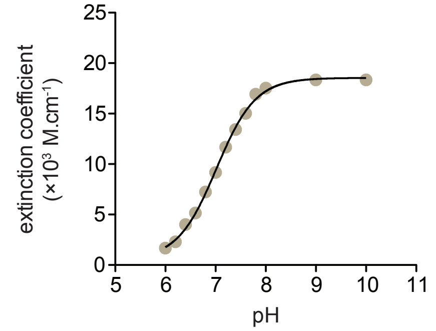


**B**

| | ε pNP (M.cm⁻¹) |
|---|---|
| pH = 7.5 | 14522 |
| pH = 8.5 | 18156 |
| pH = 9.5 | 18512 |
| pH = 10.5 | 18534 |

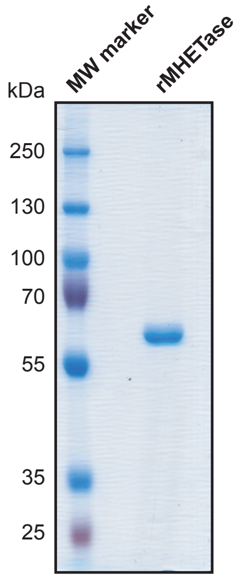


**Figure S2. para-nitrophenol extinction coefficient modeling. A.** Extinction coefficient curve fitting. Discreet extinction coefficient data from Biggs 1954 [46] was used to model para-nitrophenol extinction coefficients between pH 6 and 10. **B.** Extinction coefficients for para-nitrophenol at the indicated pH used in this study based on modelling shown in A.

**Figure S3. Purified recombinant MHETase.** Molecular weights of reference markers in kDa are indicated
