## Supplemental Tables for "Development of a yeast whole-cell biocatalyst for MHET conversion into terephthalic acid and ethylene glycol"

### Table S1. Yeast strains used in this study.

| strain ID | shorthand | genotype |
|---|---|---|
| DHY213 | - | *MAT***a** *CAT5(91M) SAL1 MIP1(661T) HAP1 MKT1(30G) RME1(INS-308A) TAO3(1493Q) leu2Δ0 his3Δ1 ura3Δ0 met15Δ0* |
| RLKY218 | - | *MAT***a** *CAT5(91M) SAL1 MIP1(661T) HAP1 MKT1(30G) RME1(INS-308A) TAO3(1493Q) leu2Δ0 his3Δ1 ura3Δ0 met15Δ0 leu2Δ::RNR2pr-tetR-NLS-tup1-7tet.1pr-tetR-NLS[LEU2]* |
| RLKY229 | SED1 | *RLKY218 can1Δ::WTC846pr-OST1-GFP-TEV-myc-SED1-tPRM9* |
| RLKY234 | AGA2 | *RLKY218 can1Δ::WTC846pr-OST1-GFP-TEV-myc-AGA2-tPRM9* |
| RLKY230 | CCW12 | *RLKY218 can1Δ::WTC846pr-OST1-GFP-TEV-myc-CCW12-tPRM9* |
| RLKY235 | CIS3 | *RLKY218 can1Δ::WTC846pr-OST1-GFP-TEV-myc-CIS3-tPRM9* |
| RLKY231 | TIP1 | *RLKY218 can1Δ::WTC846pr-OST1-GFP-TEV-myc-TIP1-tPRM9* |
| RLKY232 | CWP2 | *RLKY218 can1Δ::WTC846pr-OST1-GFP-TEV-myc-CWP2-tPRM9* |
| RLKY224 | M-AGA2 | *RLKY218 can1Δ::WTC846pr-OST1-MHETase-GFP-TEV-myc-AGA2-tPRM9* |
| RLKY225 | M-CCW12 | *RLKY218 can1Δ::WTC846pr-OST1-MHETase-GFP-TEV-myc-CCW12-tPRM9* |
| RLKY226 | M-CIS3 | *RLKY218 can1Δ::WTC846pr-OST1-MHETase-GFP-TEV-myc-CIS3-tPRM9* |
| RLKY227 | M-TIP1 | *RLKY218 can1Δ::WTC846pr-OST1-MHETase-GFP-TEV-myc-TIP1-tPRM9* |
| RLKY228 | M-CWP2 | *RLKY218 can1Δ::WTC846pr-OST1-MHETase-GFP-TEV-myc-CWP2-tPRM9* |
| RLKY244 | M-SED1 | *RLKY218 can1Δ::WTC846pr-OST1-MHETase-GFP-TEV-myc-SED1-tPRM9* |
| RLKY245 | intra-M | *RLKY218 can1Δ::WTC846pr-MHETase-GFP-TEV-myc-tPRM9* |
| RLKY247 | secretion-M | *RLKY218 can1Δ::WTC846pr-OST1-MHETase-GFP-TEV-myc-tPRM9* |
| GFP library strains | | *MAT***a** *xxx::GFP[HIS3MX] leu2Δ0 his3Δ1 ura3Δ0 met15Δ0* |

**Table S2. DNA sequences of the MHETase surface display chimeras and display partners**

**MHETase chimera sequences:**

acgccgccatccagtgtttaaacgaactagtgcggccgccagttcgagtttatcattatcaata
ctgccatttcaaagaatacgtaaataattaatagtagtgattttcctaactttatttagtcaaa
aaattagcctctatcattgatagagtgtctgggtgatctatcattgatagagcatccactaatt
ttaattctgctgtaacccgtacatgcccaaaatagggggcgggttacacagaatctatcattga
tagagtgtctgggtgatctatcattgatagagcatccactaaatataatggagctctatcattg
atagagcatccactaaaaaaaagaatcccagcaccaaaatattgttttcttcaccaaccatcag
ttcataggtccattctcttagcgcaactacagagaacaggggcacaaacaggcaaaaaacgggc
acaacctcaatggagtgatgcaacctgcctggagtaaatgatgacacaaggcaattgacccacg
catgtatctatctcattttcttacaccttctattaccttctgctctctctgatttggaaaaagc
tgaaaaaaaaggttgaaaccagttccctgaaattattcccctatctatcattgatagatataaa
tatctatcattgatagagtaattctgtaaatctatttcttaaacttcttaaattctactttat
agttagtctttttttagttttaaaacaccaagaacttagtttcgaataaacacacataaaca
aaATGCGTCAAGTCTGGTTTTCTTGGATTGTTGGATTGTTTTTATGCTTTTTTAATGTTTCATC
GGCAGGTGGCGGAAGCACACCCCTACCCCTACCACAACAACAACCGCCCCAGCAGGAGCCCCCT
CCACCACCGGTACCACTAGCTTCAAGGGCGGCATGTGAAGCTTTGAAAGATGGCAATGGAGACA
TGGTTTGGCCGAATGCTGCTACTGTGGTTGAGGTGGCTGCATGGCGTGACGCCGCGCCCGCTAC
TGCAAGTGCGGCGGCTCTGCCTGAGCATTGTGAAGTATCAGGTGCTATAGCGAAACGTACTGGA
ATCGACGGATATCCCTATGAAATTAAGTTTCGTCTACGTATGCCAGCTGAATGGAATGGCCGTT
TCTTTATGGAAGGTGGGAGTGGTACGAATGGCAGCCTAAGTGCGGCAACCGGAAGCATTGGCGG
AGGGCAGATTGCCTCCGCTCTATCCAGGAACTTTGCAACGATAGCTACGGACGGTGGCCATGAT
AACGCTGTAAATGACAATCCCGACGCCTTGGGCACAGTAGCATTCGGATTAGATCCCCAGGCAA
GACTTGATATGGGGTATAATAGTTACGACCAAGTAACTCAAGCAGGCAAGGCAGCGGTTGCCCG
TTTCTACGGTAGAGCAGCAGACAAAAGTTATTTCATCGGGTGTTCTGAAGGTGGGAGAGAAGGC
ATGATGCTTTCACAAAGATTCCCGTCTCACTACGACGGTATTGTAGCGGGCGCGCCAGGTTACC
AACTACCGAAAGCTGGTATATCCGGAGCTTGGACCACACAAAGCCTAGCTCCGGCGGCGGTCGG
CCTAGATGCTCAGGGAGTACCACTAATCAACAAGTCTTTTAGTGACGCGGATTTGCATCTACTT
AGTCAGGCTATCCTTGGCACATGCGATGCTTTAGATGGCTTGGCGGATGGAATCGTGGATAACT
ATCGTGCCTGTCAAGCTGCCTTTGACCCGGCCACGGCTGCGAATCCAGCTAACGGCAGGCTCT
GCAGTGCGTGGGCGCCAAGACCGCCGATTGCTTGAGTCCCGTGCAAGTAACCGCCATCAAAAGA
GCAATGGCAGGCCCAGTAAATTCCGCAGGTACCCCGTTATATAATAGGTGGGCCTGGGACGCTG
GAATGAGCGGATTATCCGGCACAACGTATAACCAGGGGTGGCGTTCATGGTGGTTAGGTTCATT
TAATTCAAGCGCAAACAATGCTCAAAGAGTTAGTGGGTTTAGTGCCAGAAGCTGGCTTGTGGAT
TTCGCGACTCCTCCGGAACCAATGCCAATGACCCAGGTCGCCGCCAGGATGATGAAATTTGACT
TTGACATTGACCCTTTGAAGATCTGGGCAACCTCAGGCCAGTTTACCCAGAGCAGCATGGACTG
GCACGGTGCTACCTCAACTGACTTGGCTGCTTTCAGGGACAGAGGTGGCAAAATGATTCTTTAC
CATGGGATGAGTGACGCTGCCTTTAGCGCCTTAGATACTGCTGACTACTACGAGCGTCTGGGAG
CTGCAATGCCAGGGGCTGCCGGTTTTGCCCGTCTGTTTTTAGTTCCTGGTATGAACCATTGCTC
TGGCGGACCTGGAACTGATCGTTTCGATATGCTTACTCCACTTGTGGCCTGGGTGGAACGTGGC
GAGGCACCAGACCAGATCTCAGCCTGGAGCGGCACCCCAGGCTATTTCGGCGTGGCAGCAAGGA
CTAGGCCTTTGTGTCCCTACCCCCAAATCGCTAGATACAAGGGTTCTGGAGATATAAATACAGA
GGCAAACTTCGCTTGCGCCGCCCCTCCGggttctgctggttctgctgctggttctggtgaattt
ATGGTCAGTAAGGGTGAAGAATTATTCACTGGTGTTGTTCCAATCTTGGTTGAATTGGATGGTG
ATGTTAACGGTCACAAGTTTTCTGTTCGTGGTGAAGGTGAAGGTGATGCTACTAATGGTAAATT
GACCTTGAAGTTCATCTGTACCACAGGTAAATTGCCAGTTCCATGGCCAACTTTGGTTACTACT
TTGACTTATGGTGTCCAATGCTTCTCTAGATACCCAGATCATATGAAGCAACACGACTTTTTCA

AATCCGCTATGCCAGAAGGTTACGTTCAAGAAAGAACCATCTCTTTCAAGGATGACGGTACTTA
CAAAACTAGAGCCGAAGTTAAGTTCGAAGGTGATACCTTGGTTAACAGAATCGAATTGAAGGGT
ATCGACTTCAAAGAAGATGGTAACATCTTGGGTCATAAGTTGGAATACAACTTTAACTCCCACA
ACGTTTACATTACTGCCGATAAGCAAAAGAACGGTATCAAGGCTAACTTCAAGATCAGACACAA
CGTTGAAGATGGTAGTGTTCAATTGGCTGATCACTACCAACAAAACACTCCAATTGGTGATGGT
CCAGTTTTGTTGCCAGATAACCATTACTTGTCTACCCAATCTAAATTGTCTAAGGACCCAAACG
AAAAAAGAGATCACATGGTCTTGAAAGAACGTGTTACTGCTGCTGGTATTACTTTGGGTATGGA
CGAATTATACaaggaaaatttgtattttcaatctgaacaaaaattgatttctgaagaagatttg
[------------------------DISPLAY PARTNER ----------------------]
cagaagacgggagacactagcacacaactttaccaggcaaggtatttgacgctagcatgtgtcc
aattcagtgtcatttatgatttttttgtagtaggatataaatatatacagcgctccaaatagtgc
ggttgccccaaaaacaccacggaacctcatctgttctcgtactttgttgtgacaaagtagctca
ctgccttattatcacattttcattatgcaacgctttttcccgacgagagtaaatggcgaggata
cgttctctatggaggatggc

xxx: 5′ and 3′ insertion sites at CAN1
xxx: WTC846PR846 promoter
XXX: OST1 secretion signal
XXX: *Ideonella sakaiensis* MHETase
xxx: Linker
XXX: GFP
xxx: TEV protease site
xxx: myc tag
xxx: PRM9 terminator

**Display partners sequences:**
AGA2
CAGGAACTGACAACTATATGCGAGCAAATCCCCTCACCAACTTTAGAATCGACGCCGTACTCTT
TGTCAACGACTACTATTTTGGCCAACGGGAAGGCAATGCAAGGAGTTTTTGAATATTACAAATC
AGTAACGTTTGTCAGTAATTGCGGTTCTCACCCCTCAACAACTAGCAAAGGCAGCCCCATAAAC
ACACAGTATGTTTTTtga

CCW12
GCAGCAAATGTAACAACAGCCACAGTATCACAAGAGTCTACGACACTTGTTACCATCACCTCCT
GCGAAGACCATGTCTGTTCTGAGACTGTGTCTCCTGCACTGGTCTCAACTGCCACTGTGACCGT
TGATGATGTCATTACCCAGTATACAACTTGGTGTCCCTTGACAACTGAAGCTCCAAAAAACGGC
ACTTCCACCGCAGCCCCAGTGACTAGTACAGAAGCACCAAAAAATACTACCTCAGCGGCTCCGA
CGCACTCGGTTACGAGTTACACAGGTGCCGCGGCAAAGGCACTCCCTGCTGCAGGTGCTTTATT
AGCTGGAGCTGCTGCTCTATTGTTGtga

CIS3
GACGTGATCTCACAGATTGGAGATGGACAAGTGCAAGCAACTTCTGCAGCTACCGCTCAAGCCA
CTGACTCTCAAGCTCAAGCTACTACTACAGCTACCCCAACTAGTTCCGAAAAGATTAGCTCCTC
GGCATCCAAGACTTCAACCAATGCAACCTCCTCTTCCTGTGCAACGCCAAGTTTGAAGGACAGT
TCTTGTAAAAATTCTGGTACGTTAGAACTCACGCTAAAGGATGGTGTATTGACTGATGCGAAAG
GGAGAATTGGATCGATTGTTGCCAATAGGCAATTTCAGTTTGATGGCCGCCTCCACAAGCTGG
TGCTATATACGCCGCAGGTTGGTCAATTACAGAAGATGGCTACTTGGCTTTAGGCGATAGTGAC

GTTTTCTATCAGTGTCTATCAGGAAACTTTTACAACCTTTATGATCAAAATGTCGCTGAACAGT
GTAGCGCTATCCATCTGGAAGCTGTCAGCCTTGTTGATTGCtga

CWP2
GAGAGCGCCGCTGCAATATCTCAAATTACGGATGGACAGATCCAAGCGACCACAACAGCTACTA
CTGAAGCTACAACTACTGCCGCGCCATCTTCCACCGTTGAAACCGTATCCCCTAGTTCGACTGA
GACAATTTCTCAACAAACGGAAAACGGTGCTGCTAAAGCAGCTGTTGGGATGGGTGCTGGCGCA
CTTGCTGCAGCTGCAATGTTATTGtga

SED1
CAATTTTCCAACAGCACATCTGCTAGCTCTACAGACGTAACATCCAGCTCATCAATCTCGACAA
GTTCAGGTTCTGTTACAATAACGTCTTCTGAAGCGCCCGAATCCGATAACGGTACCAGTACAGC
AGCTCCAACTGAAACGAGCACGGAGGCGCCCACAACTGCCATTCCTACAAACGGTACTTCAACT
GAGGCACCAACTACCGCTATACCAACCAACGGCACCTCGACGGAAGCCCCTACTGATACAACAA
CTGAGGCACCTACGACCGCATTACCGACCAATGGTACTTCCACTGAGGCTCCTACAGACACCAC
GACAGAAGCTCCCACCACCGGATTGCCGACGAATGGAACAACTTCAGCCTTCCCCCCTACTACG
TCCTTGCCACCATCCAATACGACAACGACTCCGCCTTATAATCCATCTACAGATTATACGACAG
ACTATACAGTTGTAACTGAGTACACAACATATTGTCCTGAGCCAACCACATTCACTACTAACGG
CAAAACCTACACGGTCACTGAACCAACTACTCTAACAATCACTGATTGCCCGTGTACAATAGAA
AAACCTACTACCACGAGTACTACAGAATACACCGTAGTCACTGAATATACAACCTACTGTCCAG
AACCAACAACTTTTACCACCAATGGCAAGACCTATACTGTTACAGAACCAACAACCTTAACAAT
TACGGATTGCCCATGTACTATTGAGAAAAGTGAAGCTCCCGAATCCTCTGTTCCTGTCACAGAA
TCTAAAGGGACCACTACTAAGGAAACTGGTGTGACTACCAAGCAGACTACCGCCAATCCTTCTT
TGACTGTGTCGACTGTGGTGCCAGTTTCTTCTAGTGCATCAAGTCATTCAGTCGTTATTAATTC
AAACGGTGCTAATGTTGTCGTACCAGGAGCCTTAGGGCTGGCAGGAGTTGCTATGCTGTTTCTT
tga

TIP1
GATACTTCGGCTGCCGAAACTGCTGAGCTGCAGGCTATCATTGGGGACATTAATTCCCATTTAA
GTGATTATTTAGGTCTTGAAACTGGTAATTCTGGATTTCAAATCCCGTCAGACGTACTCAGTGT
GTATCAACAGGTGATGACTTACACAGATGATGCTTACACCACTTTGTTTTCTGAACTTGATTTC
GACGCAATAACCAAAACTATAGTCAAACTACCATGGTATACAACAAGATTATCATCAGAAATTG
CTGCTGCCTTGGCAAGCGTTTCGCCTGCTTCTTCCGAAGCAGCTAGTTCCTCGGAGGCTGCTTC
TAGTTCGAAGGCAGCGTCCTCTTCAGAAGCAACATCATCAGCTGCACCTTCATCTTCAGCCGCT
CCATCCTCTAGCGCTGCACCCTCTTCTAGTGCCGAATCATCTAGCAAGGCCGTTAGTTCCAGCG
TCGCCCCAACAACGAGTTCGGTGTCCACCTCTACGGTTGAGACCGCATCAAACGCAGGTCAAAG
GGTTAACGCAGGTGCAGCCAGCTTCGGCGCAGTTGTAGCTGGAGCGGCGGCGCTATTGTTAtga
